# Protein Tyrosine Phosphatase 4A3 (PTP4A3) modulates Src signaling in T-cell Acute Lymphoblastic Leukemia to promote leukemia migration and progression

**DOI:** 10.1101/681791

**Authors:** M Wei, MG Haney, JS Blackburn

## Abstract

T-cell Acute Lymphoblastic Leukemia (T-ALL) is an aggressive blood cancer, and currently, there are no immunotherapies or molecularly targeted therapeutics available for treatment of this malignancy. The identification and characterization of genes and pathways that drive T-ALL progression is critical for development of new therapies for T-ALL. Here, we determined that Protein Tyrosine Phosphatase 4A3 (PTP4A3) plays a critical role in disease initiation and progression by promoting cell migration in T-ALL. PTP4A3 expression was upregulated in patient T-ALL samples at both the mRNA and protein levels compared to normal lymphocytes. Inhibition of PTP4A3 function with a small molecule inhibitor and knock-down of PTP4A3 expression using short-hairpin RNA (shRNA) in human T-ALL cells significantly impeded T-ALL cell migration capacity *in vitro* and reduced their ability to engraft and proliferate *in vivo* in xenograft mouse models. Additionally, PTP4A3 overexpression in a *Myc*-induced zebrafish T-ALL model significantly accelerated disease onset and shortened the time needed for cells to enter blood circulation. Reverse phase protein array (RPPA) revealed that manipulation of PTP4A3 expression levels in T-ALL cells directly affected the SRC signaling pathway, which plays a well-characterized role in migratory behavior of several cell types. Taken together, our study revealed that PTP4A3 is a key regulator of T-ALL migration via SRC signaling, and suggests that PTP4A3 plays an important role as an oncogenic driver in T-ALL.

**Highlights:**

- A subset of T-cell Acute Lymphoblastic Leukemia (T-ALL) highly express the phosphatase PTP4A3
- PTP4A3 expression promotes leukemia development in zebrafish T-ALL models
- Loss of PTP4A3 prevents T-ALL engraftment in mouse xenograft models
- Knock-down or small molecule inhibition of PTP4A3 prevents T-ALL migration in part via modulation of SRC signaling.

## 1. Introduction

T-cell Acute Lymphoblastic Leukemia (T-ALL) is a type of aggressive hematologic malignancy, which represents 10%–15% of pediatric and 25% of adult ALL cases [1]. Despite dramatically improved treatment for ALL over the last several decades, the treatment of T-ALL lags behind the treatment of B-cell Acute Lymphoblastic Leukemia (B-ALL) and other leukemia subtypes in both immunotherapies and development of molecular targeted therapies [2]. Additionally, relapsed T-ALL remains a major clinical concern, with less than 30% of children and 10% of adults surviving relapse, and the current intensive chemotherapy regimen for T-ALL has long-term adverse side effects on patients [1, 3–5]. Therefore, more effective and selective treatment strategies, such as molecularly targeted therapies, are critically needed for T-ALL. The development of novel therapeutics requires the identification and characterization of important drivers of leukemia progression.

Protein phosphatases cooperate with kinases to precisely maintain appropriate protein phosphorylation, and have important roles in modulating the strength and duration of signaling events, making them critical for normal cellular functions. Abnormal protein phosphorylation is frequently identified in cancers and other diseases. While kinase inhibitors have achieved significant success in clinic [6], phosphatases are underexplored as drug targets, as they were historically considered “undruggable” [7, 8], largely due to the misconception that phosphatases function primarily as tumor suppressors, as well as the challenges in developing specific phosphatase inhibitors with high bioavailability. However, to date, more than 30 potential oncogenic phosphatases have been identified, and are currently being explored as drug targets in cancer therapy [9].

Protein Tyrosine Phosphatase 4A3 (PTP4A3), also known as PRL-3, is an oncogenic phosphatase that has received significant attention as a potential therapeutic target in a variety of cancers over the last decade [10]. PTP4A3 has been extensively reported as a biomarker of tumor progression and metastasis; for example, PTP4A3 expression is documented to be significantly elevated in tumor tissues compared with healthy tissues, and in advanced metastatic versus early stage tumors across a variety of solid tumor types, including breast [11], colon [12, 13], gastric [14], brain [15], and prostate [16] as well as in Acute Myelogenous Leukemia (AML) [15, 17–19]. Similarly, PTP4A3 protein was recently shown to be highly expressed in ~80% of 151 human tumor samples across 11 varied tumor types, including liver, lung, colon, breast, stomach, thyroid, pancreas, kidney, AML, bladder and prostate [20]. Elevated PTP4A3 correlates with reduced survival in patients with breast [21], gastric [22], ovarian [23], and liver [24] cancers and in AML [25, 26]. More importantly, the causative role of PTP4A3 in cancer has been functionally demonstrated by overexpression and knock-down of PTP4A3 in normal or cancer cells. For example, ectopic expression of PTP4A3 in human melanoma, breast, lung, and colorectal cancer cells has been reported to increase cell motility, migration, invasion, and proliferation *in vitro* and to accelerate tumor formation, progression, and metastasis *in vivo* [27–30]. Similarly, knock-down of PTP4A3 expression using short hairpin RNA (shRNA) led to decreased cell proliferation, adhesion, migration, and invasion in a range of solid tumors *in vitro* and inhibits primary tumor proliferation and invasion *in vivo* in colorectal, gastric and ovarian cancers and in melanoma, ultimately improving the prognosis and life span of mice [31–34]. PTP4A3 knockout also significantly suppressed tumor formation in a colon cancer mouse model induced by azoxymethane and dextran sodium sulfate [35]. Together, these data highlight the important oncogenic role of PTP4A3 in a wide variety of cancers.

As the critical role of PTP4A3 in tumor progression and metastasis has become increasingly recognized in tumors, there have been continuous efforts to develop specific PTP4A3 inhibitors, including JMS-053 [36], Compound 43 and its analogs [37], and Analog 3 [38]. While all of these small molecules target the entire PTP4A family, including PTP4A1, PTP4A2, and PTP4A3, the anti-cancer effects of these inhibitors have been demonstrated in cancer cells or xenotransplantation models. In addition, Qi Zeng’s group has successfully developed humanized PTP4A3 antibody, which can specifically target PTP4A3 expressing tumors but not PTP4A1 expressing tumors [20, 39]. These efforts support the concept that phosphatases are in fact druggable and suggest that PTP4A3 is a feasible therapeutic target in cancer.

While PTP4A3 has well-documented roles in solid tumor progression, its role in leukemia is less defined. PTP4A3 has been reported as a downstream target of FMS-like tyrosine kinase 3 internal tandem duplication (FLT3-ITD), which is a prevalent mutation in AML, where it leads to activation of activator protein 1 (AP-1) transcription factor and contributes to AML progression *in vitro* and *in vivo* via an unknown mechanism [17, 40]. Recently, it was demonstrated that PTP4A3 is highly expressed in B-cell Acute Lymphoblastic Leukemia (B-ALL) patient samples, and *in vitro* studies showed it played a role in modulating cell migration and resistance to the chemotherapy cytarabine [41]. Here we provide both *in vitro* and *in vivo* evidence for a novel role for PTP4A3 in T-ALL. We found that PTP4A3 is upregulated in primary T-ALL patient samples and in human T-ALL cell lines, and inhibition of PTP4A3 using the pan-PTP4A family inhibitor JMS-053 or PTP4A3 knock-down by shRNA resulted in decreased migratory capability of T-ALL cells, while overexpression of PTP4A3 significantly enhanced cell migration. In addition, overexpression of PTP4A3 in a Myc-induced zebrafish T-ALL model [42, 43] demonstrated that PTP4A3 facilitates T-ALL initiation and promotes early circulation of leukemic cells in the blood. Furthermore, human T-ALL cells with PTP4A3 knock-down lost the ability to engraft and circulate in mice. Reverse Phase Protein Array analysis revealed that PTP4A3 modulates the Src signaling pathway to affect cell migration in T-ALL cells. Taken together, our study identified the critical role of PTP4A3 in T-ALL onset and progression and suggests that PTP4A3 may be a targetable oncogenic driver in T-ALL.

## 2. Materials and methods

### 2.1. Antibodies, DNA plasmids, and other reagents

Antibodies used in this study, including their manufacturer, catalog number, lot number, blocking buffer used, and dilution factor are listed in Supplemental Table 1. The specificity of antibodies against PTP4A1, 2, and 3 were validated against purified protein (Supplemental Figure 1). The PTP4A inhibitor, JMS-053, was kindly provided by Dr. John S. Lazo, Elizabeth Sharlowe, and Peter Wipf (University of Virginia, Charlottesville, VA), and the Src inhibitor, SU6656, was purchased from Sigma-Aldrich (S9692).

Lentiviral packaging plasmids psPAX2 (Addgene 12260) and pMD2.G (Addgene 12259) were gifted from Didier Trono. pLenti PGK Puro DEST (w529-2) (hereafter referred to as PGK) (Addgene 19068) and pLenti PGK GFP Puro (w509-5) (Addgene 19070) were gifts from Eric Campeau & Paul Kaufman. Non-targeting control pLKO shRNA lentivirus plasmid (MISSION, SHC002) was kindly provided by Tianyan Gao and pLKO shRNAs targeting PTP4A3 were purchased from Sigma-Aldrich; target sequences are listed in Supplemental Table 2.

pENTR:*PTP4A3* (human) and pENTR:*ptp4a3* (zebrafish) Gateway Entry constructs were made by PCR amplifying *PTP4A3* and *ptp4a3* from cDNA generated from human T-ALL cells and 24-hour post-fertilization zebrafish embryos, respectively. The PCR products were subcloned into the pENTR-d-TOPO cloning vector (ThermoFisher K2400-20). The Gateway compatible zebrafish rag2 vector and generation of the *rag2:Myc* and *rag2:mCherry* construct has been previously described [42]. The *PGK:PTP4A3* and *rag2:ptp4a3* constructs were generated using the PGK destination vector and *pENTR:PTP4A3* or the rag2 destination vector and pENTR:ptp4a3 along with Gateway LR Clonase II enzyme mix, according to manufacturer’s protocol (ThermoFisher 11791020).

### 2.2. T-ALL cell lines and cell culture

All the human T-ALL cell lines used in the study were authenticated by short tandem repeat (STR) DNA profiling prior to experimentation. Cells were grown in RPMI1640 (ThermoFisher 11875119) supplemented with 10% heat-inactivated fetal bovine serum (Atlanta Biologicals, S11150H, Lot M17161). Cells were cultured at 37°C in a humidified atmosphere with 5% CO_2_.

### 2.3 Western blot

Western blot analysis was performed using a stain-free technology developed by BioRad [44, 45]. Protein was separated on TGX-stain free pre-cast 4–20% SDS gels (Biorad 4568094) at 200V for 30’. After electrophoresis, total protein was imaged using UV activation, and then transferred to polyvinylidene difluoride (PVDF) membrane. Membranes were blocked for 1hr in the appropriate blocking solution, and incubated with primary antibody as described in Supplemental Table 1. After incubation with HRP-conjugated secondary antibodies, proteins were visualized using Clarity Western ECL Substrate reaction (BioRad, 1705061), and imaged using the ChemiDoc imaging system (BioRad). The volume of the target protein bands and total protein volume of each lane were normalized and quantified using ImageLab software (Biorad). For all figures shown, a band of total protein ~50kD is size is used to represent the lane of total protein.

### 2.4. Primary human samples

Frozen isolated peripheral blood mononuclear cells (PBMCs) from T-ALL patients were kindly provided by Dr. Michelle Kelliher (University of Massachusetts). PBMCs from healthy donors were purchased from Precision for Medicine.

### 2.5. Microarray data analysis

The primary patient microarray datasets were accessed through the Gene Expression Ominbus (GEO) at NCBI (https://www.ncbi.nlm.nih.gov/geo), including GSE13159 [46, 47] and GSE14615 [48, 49].

### 2.6. Lentivirus packaging and T-ALL cells infection

For PTP4A3 knock-down, lentivirus was produced in HEK 293T cells (ATCC CRL-3216) by plating at a density of 4×10^6^ cells in a 10 cm^2^ culture dish. After 24hr, cells were transfected using 6μg of pLKO-shRNA constructs (hereafter referred as shPTP4A3 #1, 2, or 3) against PTP4A3 or the scrambled pLKO-shRNA construct (hereafter referred as SCR) in combination with 3μg psPAX2 (packaging plasmid) and 2μg pMD2.G (envelope plasmid) with 30μL *Trans*IT-LT1 transfection reagent (Mirus, MIR 2300). For T-ALL cell infection, 2.5mL virus with 10μg/mL polybrene (Thermo Fisher Scientific TR-1003-G) was added to 5×10^5^ cells plated in 6-well dishes, and centrifuged at 2250 rpm for 90 minutes. Virus was washed out with PBS after 24hr, and cells were selected in culture media with 5μg/mL puromycin for 48hr before experiments. PTP4A3 knock-down was confirmed by immunoblotting.

To generate PTP4A3-ovexpressing cell lines, 293T cells were transfected with PGK:*GFP* with or without PGK:*PTP4A3* in combination with psPAX2 (packaging plasmid) and pMD2.G (envelope plasmid) for virus packaging, as described above. T-ALL cells were infected with spin cycles and selected in medium with 5μg/mL puromycin (Jurkat) or 1μg/mL puromycin (HBPALL) for one week to produce stably expressing cell lines, then maintained in media with puromycin thereafter. PTP4A3 overexpression was confirmed by immunoblotting.

### 2.7. *In vitro* cell-based assays

The CellTiter-Glo Luminescent Cell Viability Assay (Promega, G7570) was used to measure cell survival according to the manufacturer’s instructions. A Synergy LX BioTek multi-mode plate reader was used to read luminescent signal.

To quantify cell migration, cells were first cultured in serum-free RPMI1640 overnight, then 4×10^5^ cells were seeded in triplicate wells into the upper chamber of Corning HTS Transwell 96 well permeable supports with a 5μm pore size (CLS3388-2EA). In experiments using JMS-053 or SU6656, cells were pre-treated with JMS-053, SU6656 or DMSO control for 2 hours before plating into the upper chamber. The lower chamber was loaded with 200μL of complete medium. The cells that migrated into the lower chamber were quantified by CellTiter-Glo Luminescent Cell Viability Assay.

Cell cycle was analyzed by quantifying 5’-ethynyl-2’-deoxyuridine (EdU) uptake using ClickIT EdU Alexa Fluor 647 (Thermo Fisher Scientific, C10424) according to the manufacturer’s protocol. Briefly, cells were incubated with Edu for 2 hrs., and then fixed in 4% paraformaldehyde before Alexa Fluor conjugation. DAPI (0.1μg/ml) (ThermoFisher 62248) was also used to stain the DNA.

Apoptosis was quantified by staining cells with Annexin V APC (ThermoFisher 88-8007-74) according to the manufacturers protocol in the presence of DAPI (0.05μg/ml).

### 2.7. RPPA assay and data processing

Reverse Phase Protein Array (RPPA) and data analysis were performed by the RPPA Core Facility at MD Anderson Cancer Center as previously described [50]. Briefly, total protein extracted from cells was printed on nitrocellulose-coated slides using an Aushon Biosystems 2470 Arrayer. Slides were probed with antibodies previously validated by the core, and signals reflecting protein abundance were visualized by a secondary streptavidin-conjugated HRP and DAB colorimetric reaction. The slides were scanned, analyzed, and quantified using Array-Pro Analyzer software (MediaCybernetics) to generate spot intensity. SuperCurve GUI was used to estimate relative protein levels (in log2 scale). A fitted curve (“supercurve”) was created with signal intensities on the Y-axis and relative log2 amounts of each protein on the X-axis using a non-parametric, monotone increasing B-spline model. Raw spot intensity data were adjusted to correct spatial bias before model fitting using “control spots” arrayed across the slide. A QC metric was generated for each slide to determine slide quality and only slides with 0.8 on a 0-1 scale were used. Protein measurements were corrected for loading using median-centering across antibodies. Samples with low protein levels were excluded from further analysis.

### 2.9. Zebrafish T-ALL models

Use of zebrafish was approved by the University of Kentucky’s Institutional Animal Care and Use Committee (IACUC), protocol 2015-2225 and all experiments were performed in accordance with their guidelines. Microinjections of 15ng/μL *rag2:Myc* + 45ng/μL *rag2:mCherry* or 15ng/μL *rag2:Myc* + 15ng/μL *rag2:ptp4a3* + 30ng/μL *rag2:mCherry* were used to generate zebrafish T-ALL in CG1 strain zebrafish as previously described [43, 51]. Zebrafish were monitored for leukemia onset and progression starting at 21 days post-fertilization and every 3 days onwards by analyzing the percent of the body expressing mCherry-positive leukemia cells using a Nikon fluorescence-equipped SMZ25 microscope. Circulating mCherry-positive T-ALL was noted by examining the vessels within the tail vasculature. Animals were monitored until 90 days post-fertilization or until they had to be sacrificed due to leukemia burden.

Zebrafish tumors were harvested by maceration with a razor blade in 1-4 mL. 0.9x PBS with 5% Fetal Bovine Serum when > 70% of the animal was overtaken by leukemia cells (J.S. Blackburn, Liu, Sali, Langenau David M., 2011), passed through a 40 μm mesh strainer, and fluorescent leukemia cells were counted on Countess II FL cell counter (ThermoFisher Scientific). 5-8×10^4^ cells in 100 μL of 0.9x PBS were loaded into EZ Single Cytofunnel (ThermoFisher Scientific A78710003) containing coated Shandon Cytoslides (ThermoFisher Scientific 5991056) and spun at 500 rpm for 5 minutes on Shandon Cytospin 3. Slides were allowed to dry overnight before staining. For May-Grunwald Giemsa staining, slides were incubated in 1:2 dilution of May-Grunwald stain (Sigma MG500-500ML) in distilled water for 5 minutes. Slides were then moved into a 1:10 dilution of Giemsa stain (Sigma GS500-500ML) in distilled water for 30 minutes and washed in distilled water before imaging on a BioTek Lionheart FX microscope.

To assess gene expression, zebrafish leukemias were harvested as previously described [43], and RNA was isolated from the cells using Zymo Research Quick-RNA kit (R1054). Total RNA was reverse transcribed (BioRad iSCRIPT, 1708891) and real time PCR performed using iTaq Universal Sybr Green Supermix (Biorad, 1725120), primer sequences available in Supplemental Table 3. Data were normalized to ef1a expression and fold change was calculated using the 2^−ΔΔ^Cq method.

### 2.10. Xenograft models in immune-compromised mice

Use of mice was approved by the University of Kentucky’s IACUC, protocol 2017-2754. Eight-week old NOD.Cg-Prkdc^*scid*^Il2rg^tm1Wjl^/SzJ (NSG) mice were obtained from Jackson Laboratory. The mice were randomized by placing into groups such that the difference between average group weight is not greater than 10%. Jurkat cells were infected with Scrambled shRNA or PTP4A3 shRNA as described above. Two days after virus infection, Jurkat cells were selected using 5μg/ml puromycin for two days, stained with trypan blue, and viable cells were FACS isolated. 10^6^ live cells in 100μL PBS were injected intravenously. Peripheral blood samples (100-150μL) were collected by submandibular bleeding at 4, 6 and 8 weeks post-transplantation and stained with human CD45 antibody according to manufacturer’s protocol and analyzed by flow cytometry.

### 2.11. Statistical analysis

Results are shown as mean ± standard deviation. Statistical analyses were performed using GraphPad Prism 7. Two-tailed t-tests were performed to compare two groups with similar distribution, and Analysis of Variance [52] with Tukey’s multiple comparisons was used to compare more than two groups. Human microarray data were analyzed using two-sample t-test and Wilcoxon rank sum tests, and survival curves were analyzed using Log-rank tests. Difference in circulating zebrafish cells were analyzed using two sample t-test between percents. All bar graphs shown are data pooled from ≥3 experiments.

## 3. Results

### 3.1. PTP4A3 is highly expressed in T-ALL patient samples and T-ALL cell lines

Given the important role of PTP4A3 in solid tumor progression and its emerging role in some types of leukemia, we wanted to define the function of PTP4A3 in T-ALL. We first investigated PTP4A1-3 expression in ALL patients and healthy peripheral blood mononulcear cells (PBMCs) using gene expression datasets deposited in the Oncomine database (GSE13159). Data revealed that PTP4A3 mRNA expression is significantly higher (*p*=6.8e-10) in T-ALL samples (n=174) compared to that in PBMCs (n=72, Figure 1A). In GSE14615, we saw no significant difference in *PTP4A3* expression between patients who achieved remission (n=29) versus those that relapsed (n=11) or suffered induction failure (neen patients who achieved remission (n=29) versus those that relapsed (n=11) or suffered induction failure (n=7, Figure 1B). However, the sample size was relatively small in this study, and further investigation is required to determine whether PTP4A3 expression is a predictor for T-ALL treatment failure. We also investigated the expression of PTP4A1 and PTP4A2, two closely related phosphatases in the PTP4A family. PTP4A1 expression is significantly lower in T-ALL patient samples compared to PBMCs, while PTP4A2 expression is significantly higher. Neither PTP4A1 nor PTP4A2 are significantly associated with disease relapse (Supplementary Figure 2).

**Figure 1.**
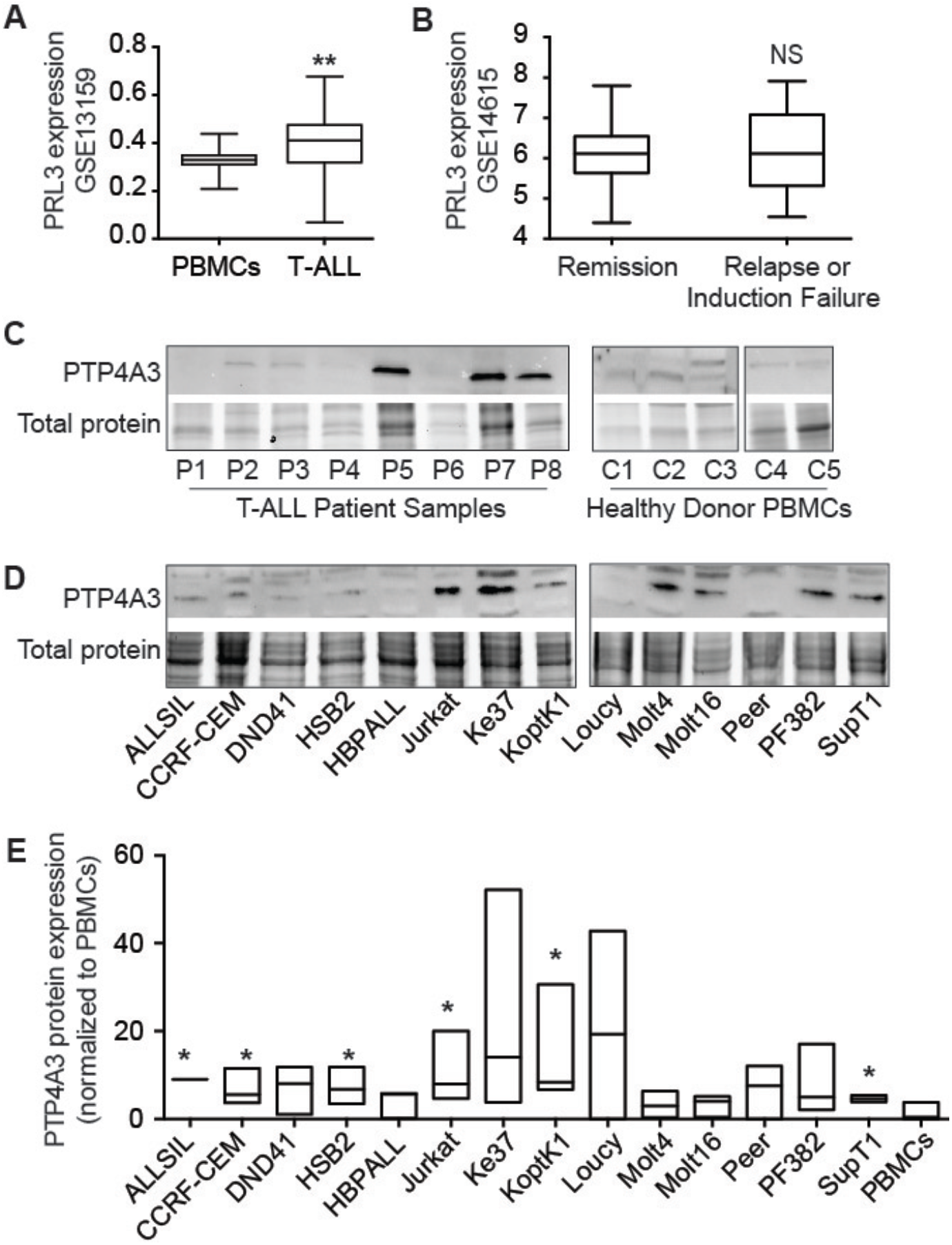
PTP4A3 is highly expressed in a majority of human T-ALL. (A) Microarray expression analysis of GSE13159 comparing normal peripheral blood mononuclear cells (PBMCs) and primary T-ALL, \*\**p*=6.8e-10. (B) Analysis of GSE14615 comparing PTP4A3 expression between samples from T-ALL patients achieving remission and the patients with induction failure. *NS*=not significant. Representative western blot analysis of (C) primary patient T-ALL and PBMCs, and (D) human T-ALL cell lines, showing PTP4A3 expression, as well as a subset of the total protein loaded. E) Quantification of 3 independent western blots across cell lines, showing PTP4A3 expression can vary, even within the same cell line. \**p*<0.05, compared to PBMCs.

We next assessed PTP4A3 protein expression in T-ALL patient samples. Western blot analysis showed approximately 50% of patient T-ALL samples expressed PTP4A3 protein, while it is detected at only low levels, if at all, in PBMCs from 5 healthy donors (Figure 1C).

Finally, we assessed PTP4A3 protein expression across 14 human T-ALL cell lines, which expressed PTP4A3 protein at varying levels (Figure 1D-E), with a majority of cell lines having higher PTP4A3 expression than normal PBMCs. Consistent with gene expression datasets, PTP4A1 protein was not detected across T-ALL cell lines but PTP4A2 is expressed at high levels in most of the T-ALL cell lines examined (Supplementary Figure 3).

### 3.2 PRL inhibitor, JMS-053, inhibits human ALL growth and migration *in vitro*

PTP4A3 knock-down or inhibition has been shown to impair cell growth in ovarian and gastric cancer cells [33, 53]. To define the functional effects of PTP4A3 expression in T-ALL, we examined the effects of small molecule inhibition, using the non-competitive PTP4A inhibitor, JMS-053 [36] on T-ALL cell lines. Cell viability analysis showed that JMS-053 inhibited T-ALL viability after 72hr treatment at a relatively low concentration, 3-4 μM (Figure 2A), with lesser to no effects in cell lines that did not routinely express high levels of PTP4A3. Although PTP4A inhibition increased apoptosis in T-ALL cells after 24hr, the effects were not significant across multiple experiments (Figure 2B), and JMS-053 treatment had no effect on cell cycle, measured by EdU uptake (Figure 2C).

**Figure 2.**
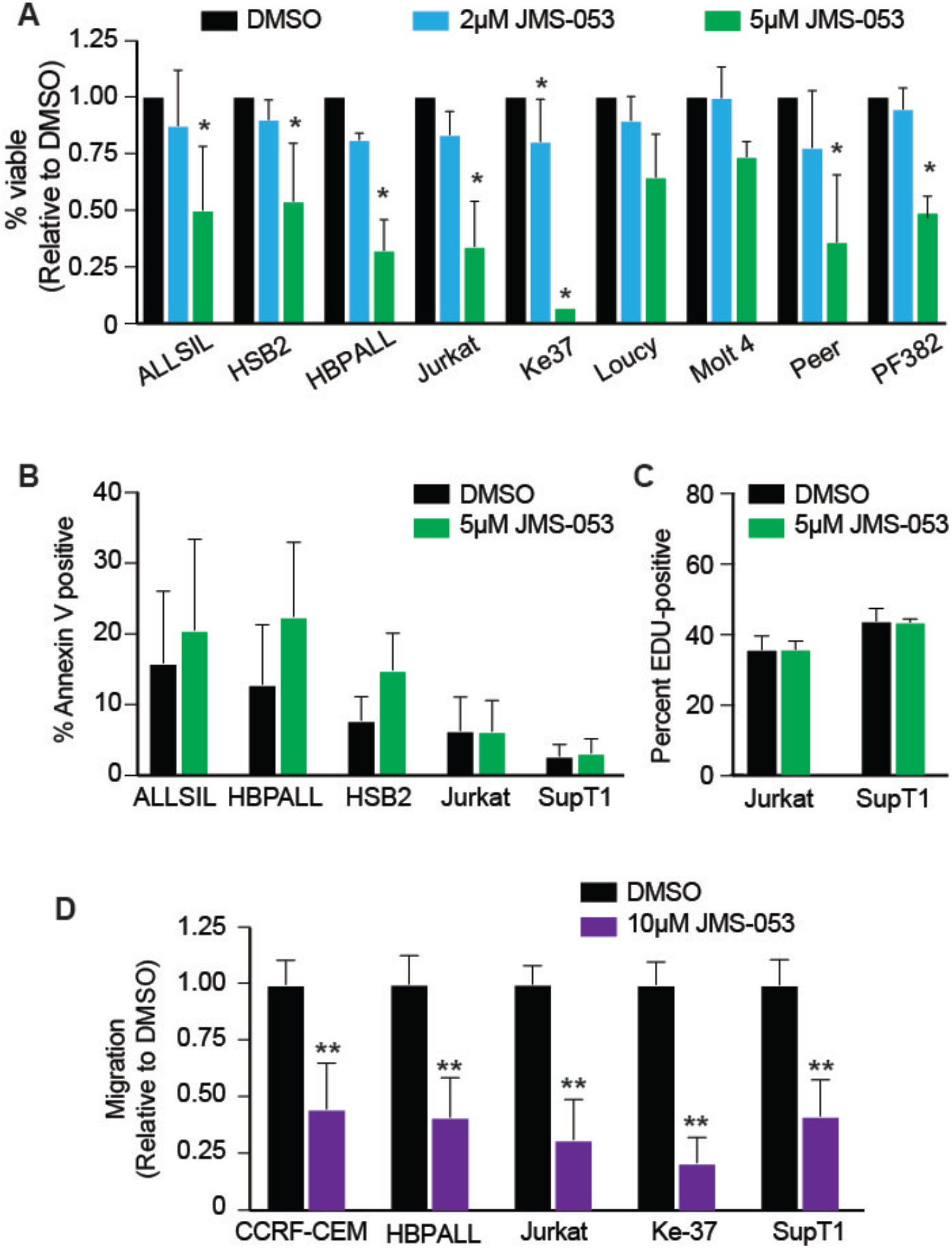
The specific pan-PTP4A inhibitor JMS-053 inhibits cell growth and migration of T-ALL cells. (A) JMS-053 inhibits cell viability, evaluated by quantifying ATP production via Cell-Titer Glo, \**p*≤0.001 or NS=not significant, compared to DMSO. There is no significant difference in apoptosis (B) or cell cycle (C) in JMS-053 treated cells, compared to DMSO control. (D) JMS-053 treatment (10μM) for 2hr suppressed cell migration of T-ALL cells, \*\**p*<0.001 compared to DMSO. For all, bars are the average of three experiments, each done in triplicate, ± standard deviation.

Previous studies have shown that PTP4A3 is also involved in migration of cancer cells [32, 34], leading us to examine whether JMS-053 affects the capability of T-ALL cells to migrate towards a serum stimulus. A panel of T-ALL cell lines that express high levels of PTP4A3 were cultured in serum free medium overnight, treated with 10μM JMS-053 for 2hrs, and plated in the upper chamber of a trans-well support. Media supplemented with 10% FBS was used as a chemoattractant in the lower chamber. After 4hrs, we quantified the migrated cells and found that JMS-053 treatment significantly (*p*<0.001) impaired the migration capability of all T-ALL cell lines tested, reducing cell migration by 30-80% (Figure 2D). Cell viability was not affected during the short time course of these experiments. These results demonstrate that PTP4A inhibition not only decreases cell viability, but also inhibits the migratory ability of T-ALL cells *in vitro*.

### 3.3 PTP4A3 knock-down in T-ALL cell lines inhibits cell migration while PTP4A3 overexpression promotes cell migration

Although JMS-053 shows high specificity towards PTP4A3 over a panel of other protein phosphatases, it also strongly inhibits PTP4A1 and PTP4A2 [36]. The phenotypes we observed in T-ALL cells treated with JMS-053, including the inhibition of cell growth and migration, could be attributed to the effects of inhibition of both PTP4A3 and PTP4A2, as PTP4A2 is also highly expressed in these cell lines (Supplemental Figure 3). In order to investigate the exact role of PTP4A3 in T-ALL, we knocked down its expression in Jurkat cells, which have high endogenous PTP4A3 expression, using shRNA. Western blot analysis showed that, 4 days after lentiviral infection, PTP4A3 was knocked-down completely by shRNA constructs #2 and #3 compared to scrambled (SCR) control shRNA (Figure 3A). We also probed for PTP4A2 expression in the PTP4A3 knock-down cells to determine if there might be any compensation from closely related PTP4A proteins, and found that PTP4A2 expression was not affected by PTP4A3 knock-down (Supplemental Figure 4A). Interestingly, silencing PTP4A3 did not last long enough to generate stable cell lines. We found that despite puromycin selection, PTP4A3 expression levels return over time (Supplemental Figure 4B). These data suggest that cells with higher PTP4A3 expression may outcompete those with stronger knock-down. We therefore performed transient knock-down of PTP4A3 instead of generating stable cell lines to complete all the PTP4A3 shRNA related experiments.

**Figure 3.**
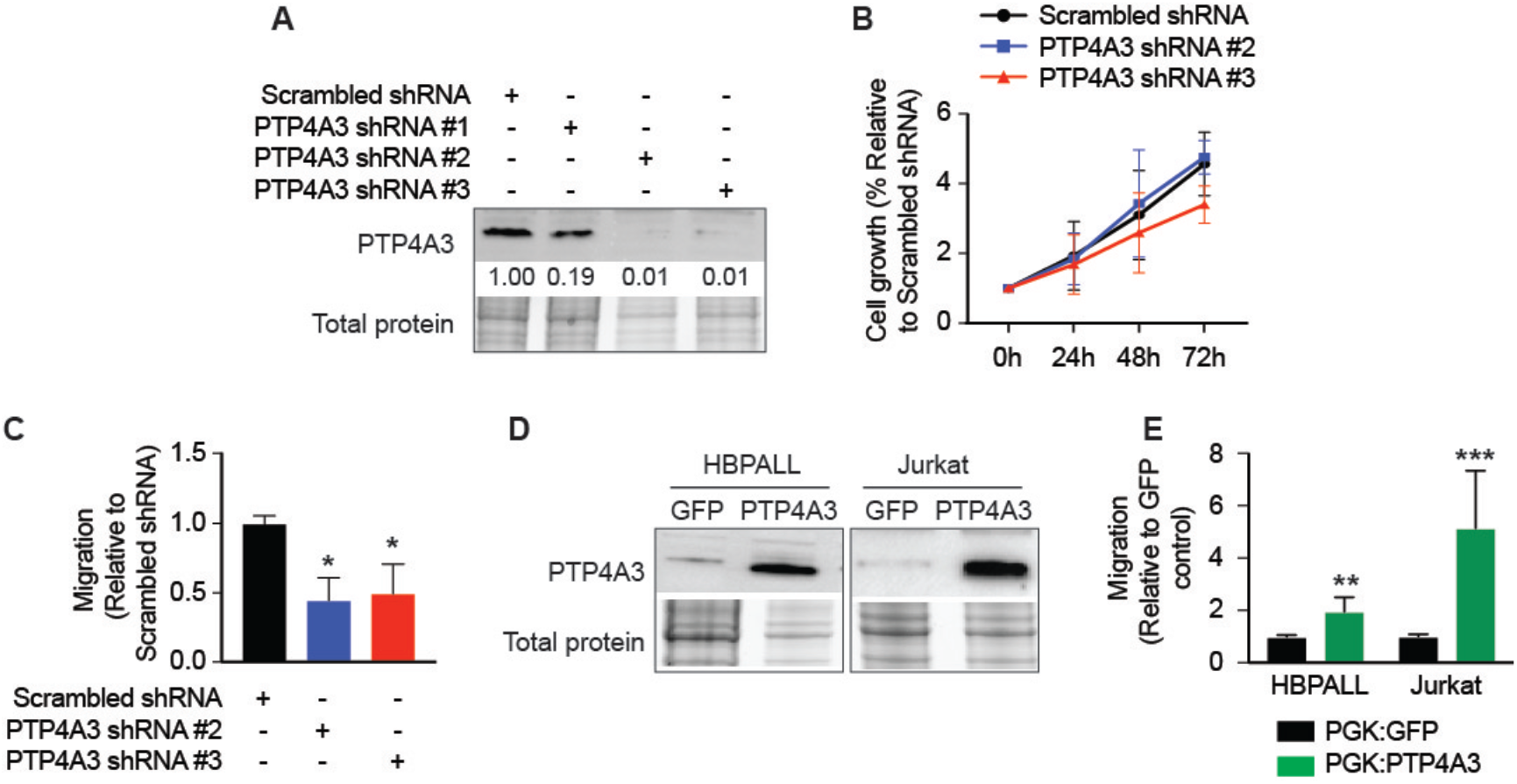
PTP4A3 promotes cell migration, but not growth, in T-ALL. (A) Representative western analysis figure showing PTP4A3 protein expression in Jurkat T-ALL cells 4d post-infection with lentivirus carrying shRNA. Numbers represent relative expression of PTP4A3 protein, normalized total protein and compared to scrambled (SCR) control. (B) Cells infected with SCR or PTP4A3 knock-down shRNA were cultured in media 4 days post-infection with 5μg/ml puromycin for 72 additional hours. Cell growth was determined by Cell Titer-Glo assay and normalized to the readout of day 0, and show no difference between groups. Data shown are the average of 3 independent experiments, done in triplicate, *NS*=not significant. (C) Knock-down of PTP4A3 in Jurkat cell line reduced migration towards a serum stimulus more than 50%. Migration was normalized to the cells infected by SCR shRNA, p<0.05 compared to SCR control. (D) Western blot analysis showed expression of PTP4A3 in Jurkat and HBP-ALL cell lines after infection with lentivirus harboring PGK:GFP or PGK:PTP4A3. (E) High PTP4A3 expression increased cell migration in T-ALL cells. Migration was normalized to control cells expressing GFP only, \*\**p*<0.001, \*\*\**p*<0.0001. For all bar graphs, data are the average of at least three independent experiments, done in triplicate.

Proliferation of PTP4A3 knock-down Jurkat cells was comparable to SCR control cells, indicating that knock-down of PTP4A3 did not affect cell viability over time (Figure 3B). However, silencing PTP4A3 expression significantly reduced cell migration by ~50% (*p*<0.02, Figure 3C). Similarly, when we overexpressed PTP4A3 in Jurkat and HBPALL cells (Figure 3D), we found no significant difference in cell growth (Supplemental Figure 5), but significantly increased migratory capability compared to control (*p*≤0.009, Figure 3E). Together, these data suggest that PTP4A3 plays an important role in regulating cell migration, but not proliferation, in T-ALL cells.

### 3.4 PTP4A3 increases circulation of T-ALL cells in a zebrafish model

Although the oncogenic role of PTP4A3 has been reported in solid tumors, its functional role in T-ALL progression has not been demonstrated [10]. The high expression of PTP4A3 in T-ALL patient samples and its role in promoting migration in T-ALL cell lines suggests it may play an oncogenic role in T-ALL. We used a zebrafish *Myc*-induced T-ALL model [42, 51] to assess the role of PTP4A3 in T-ALL onset and progression. One-cell stage zebrafish embryos were injected with plasmids containing *rag2:Myc* with *rag2:mCherry*, and with or without *rag2:ptp4a3*. T-ALL develops in zebrafish from the thymus and expands into local tissues before entering the circulation. Fish were monitored for leukemia growth by quantifying the percent mCherry-positive cells within the body of the animal; >70% mCherry-positive was considered leukemic. Zebrafish *ptp4a3* has 88% homology to human *PTP4A3* with conservation of the critical domains [54]. Zebrafish T-ALL that expressed *ptp4a3* expanded from the thymus earlier (Figure 4A) than T-ALLs expressing *Myc* alone, and overall, animals with *Myc*+*ptp4a3* T-ALL were leukemic approximately 10 days faster than *Myc* control animals, indicating Ptp4a3 enhances T-ALL onset (Figure 4B). The lymphoblasts were morphologically similar between the groups (Figure 4D), and gene expression analyses indicated that both the *rag2:Myc* and *rag2:Myc*+*rag2:ptp4a3* leukemias expressed the lymphocyte specific genes *rag1* and *rag2* and the T-cell genes *lck* and *tcrB*, but not B-cell related genes *igD* or *igM*, indicating all leukemias generated were of T-cell origin. We were also able to verify that the *rag2:Myc*+*rag2:ptp4a3* leukemias expressed >10-fold higher levels of Ptp4a3 than the *Myc* control group (Figure 4C). Interestingly, endogenous *ptp4a3* expression was also significantly higher in the *rag2:myc* T-ALLs than normal blood, suggesting that ptp4a3 may be an important collaborating oncogene in T-ALL development.

**Figure 4:**
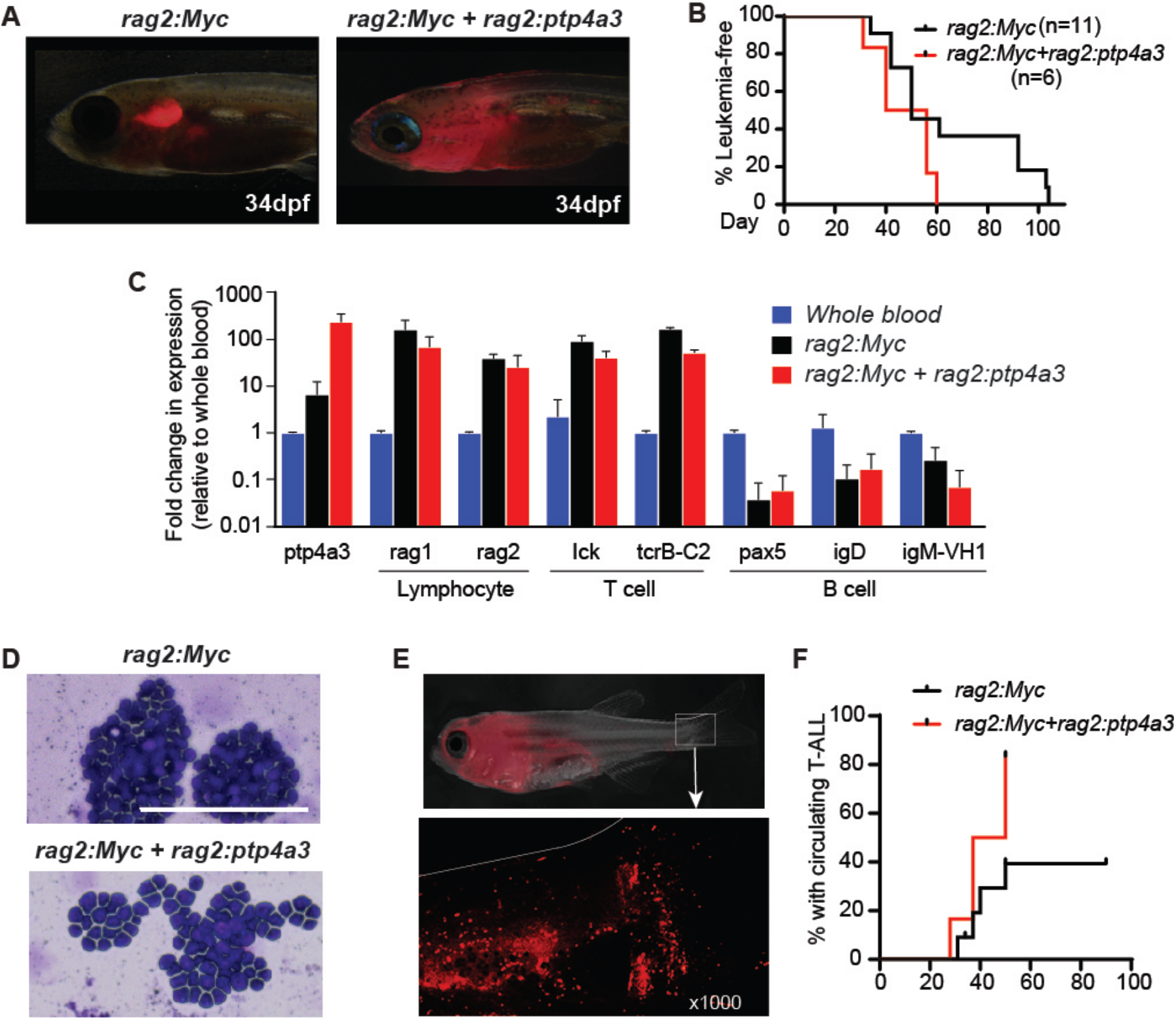
PTP4A3 enhances circulation of leukemia cells in a zebrafish T-ALL model. (A) Representative images of transient transgenic zebrafish expressing *rag2:Myc*+*rag2:mCherry* (n=11) or *rag2:Myc*+*rag2:mCherry*+*rag2:ptp4a3* (n=6) at 34 days post-fertilization (dpf). (B) Kaplan-Meier analysis of time (days) to leukemia (>70% of animal is mCherry-positive), showing PTP4A3 expression enhances T-ALL progression. (C) Representative images of May-Gunwald Giemsa staining of blood samples from fish from each leukemia type. Scale bar=100μm. (D) Realtime RT-PCR analysis of lymphocyte, T-cell, and B-cell specific genes. Bars are the average expression of three samples per group. (E) Representative *rag2:Myc*+*rag2:mCherry*+*rag2:PTP4A3* animal, showing circulating mCherry+ leukemia cells within the tail fin. (F) Kaplan-Meier analysis of time (days) for each T-ALL to be visualized in circulation.

Because the T-ALL cells were fluorescently labeled, we were also able to determine the time at which the T-ALL cells begin to circulate by visualizing cells within the vasculature in the tail fin (Figure 4E, Supplemental Videos 1 and 2). While less than half of zebrafish with *Myc*-expressing T-ALL had circulating cells when >50% of the animal was mCherry-positive, more than 80% of the *Myc*+*ptp4a3* expressing T-ALLs were circulating at a median time point of 42d (Figure 4F). Taken together, these data suggest that PTP4A3 can play an important role in T-ALL onset and progression, likely by enhancing migration into local tissues and contributing to the ability of the cells to enter circulation.

### 3.5 PTP4A3 knock-down impairs human T-ALL engraftment *in vivo*

In order to determine whether silencing PTP4A3 expression in human T-ALL affects their oncogenic ability *in vivo*, xenotransplantation of T-ALL was performed using the PTP4A3 knock-down Jurkat cells. As shown in Figure 5A, Jurkat cells expressing a scrambled shRNA or an shRNA targeting *PTP4A3* were injected intravenously into immune-compromised mice. At 4, 6, and 8 weeks after transplantation, blood samples were collected and stained with anti-human CD45. Flow cytometry showed no human CD45-positive cells in the circulation of mice injected with PTP4A3 knock-down Jurkat cells, while mice injected with Jurkat expressing scrambled shRNA cells showed increasing numbers of CD45 positive cells in the blood each week (Figure 5B). Three mice harboring scrambled shRNA expressing T-ALL had to be euthanized before the 8-week time point due to mobility issues likely resulting from T-ALL infiltration into the spinal column. Mice with PTP4A3 knock-down remained healthy throughout the study. Together, these data show that PTP4A3 significantly affects engraftment and circulation of human T-ALL cells xenografted into mice.

**Figure 5.**
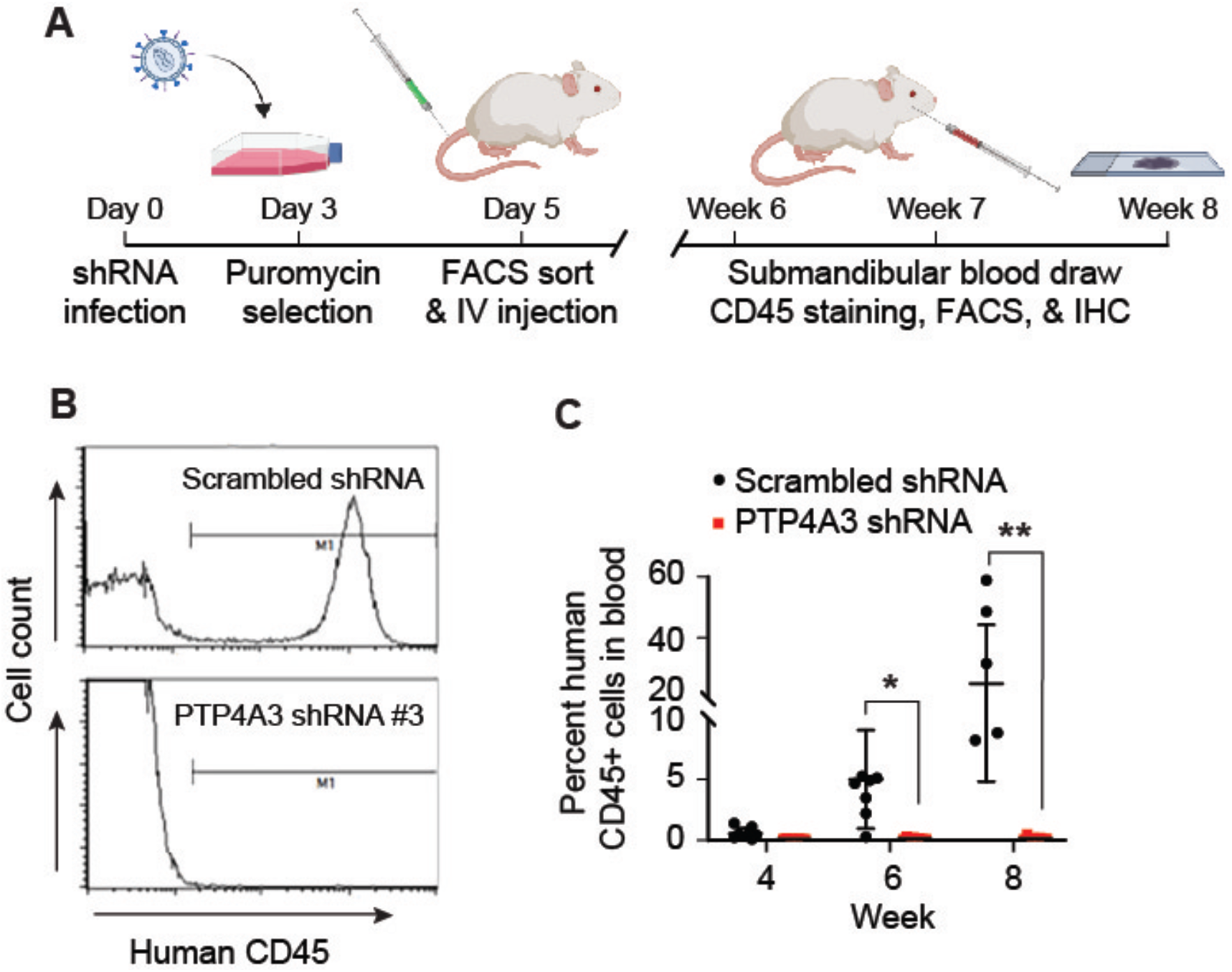
PTP4A3 knock-down impairs T-ALL engraftment in a xenograft mouse model. (A) Schematic diagram of the xenotransplantation assay. (B) PTP4A3 knocked-down Jurkat cells were not detected in transplanted mice. Left: Representative flow cytometry analysis of submandibular blood sample after human CD45 staining. Right: Quantification of human CD45 staining of mice blood at week 4,6 and 8 after transplantation. Each dot represents one mouse, the horizontal line represents the mean value, and the standard deviation is shown, *p<0.01 and **p<0.001 compared to shRNA control xenografted mice.

### 3.6 PTP4A3 inhibition inhibits SRC pathway signaling to decrease migration

Our data suggest that PTP4A3 functions in T-ALL progression by modulating leukemia cell migration. To determine the mechanism by which PTP4A3 might contribute to cell motility, we analyzed the effect of PTP4A3 expression on 442 different proteins, including 86 phospho-proteins using Reverse-Phase Protein Array (RPPA). Total protein from PTP4A3 knock-down or control Jurkat cells was extracted and RPPA assays were performed using validated antibodies. After rigorous quality control for data analysis, we identified 23 proteins that showed greater than 20% differential expression between scrambled PTP4A3 shRNA or PTP4A3 knock-down T-ALL cells (Figure 6A and Supplemental Table 4). RPPA on PTP4A3 overexpressing compared to control cells identified 21 proteins with more than 30% differential expression (Figure 6B and Supplemental Table 5). We found that phosphorylation of tyrosine residue 527 of Src (Src_pY527) was increased in PTP4A3 knock-down cells, and decreased in cells with PTP4A3 over-expression.

**Figure 6.**
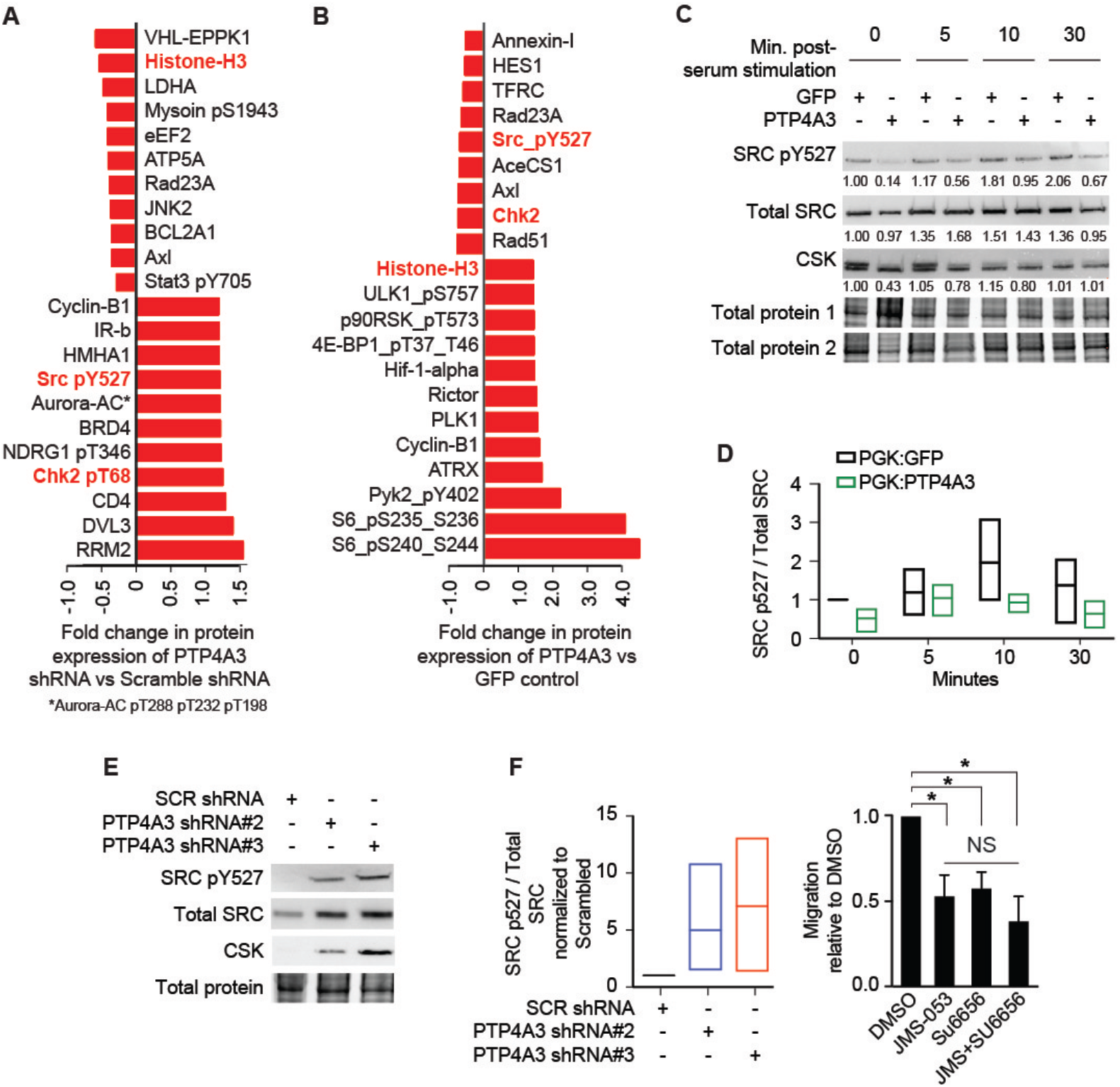
PTP4A3 modulates Src phosphorylation to regulate cell migration. Reverse phase protein array analysis (RPPA) of (A) PTP4A3 knock-down or (B) over-expression of PTP4A3 in Jurkat cells showed differential protein expression when compared to controls. Red bars show any protein that was up or down regulated 20%, and protein names shown in red are common in both groups, and include Chk2, Histone H3, and Src_pY527. (C) Western blot validation of Src pathway in PTP4A3 overexpressing Jurkat cells. Cells were serum starved overnight and added to serum containing complete media for the indicated time points. Src_pY527 and CSK protein were consistently decreased in PTP4A3 over-expressing Jurkat cells but not GFP expressing control cells. Numbers shown represent relative protein expression. (D) Quantification of Src_pY527 western blots representing at least n=3 independent experiments. Box plots show the mean and standard deviation. (E) Representative western blot analysis of Src_pY527, total Src, and CSK in Jurkat cells with PTP4A3 knock-down, and (F) quantification of at least 3 independent experiments, showing the mean and standard deviation of each group. (G) Cells with PTP4A3 over-expression were treated with DMSO, Src inhibitor Su6656 (2.5μM), JMS-053 (10μM) or in combination, and placed in a trans-well migration assay towards a serum stimulus. Migrated cells were quantified using Cell-Titer glo, \**p*<0.01 compared to DMSO and NS=not significant compared to JMS-053 treatment alone.

The activation of Src, a non-receptor kinase, occurs in a large fraction of cancers and has a prominent role in cell migration and metastasis [55]. Its activity is regulated by phosphorylation of tyrosine 527, which is an inhibitory phosphorylation site and also regulated by the CSK (C-terminal Src Kinase) and various other phosphatases. We further validated the changes in Src pathway expression seen in RPPA studies using western blot in PTP4A3 knock-down or overexpressing T-ALL cell lines. PTP4A3 overexpressing and control Jurkat cells were serum-starved overnight, then placed in complete media for 0, 5, 15, and 30 minutes. Cells with PTP4A3 overexpression had decreased phosphorylation of Src_p527 at all time points compared to control (Figure 6C-D), while total Src protein expression was not affected. Interestingly, the overexpression of PTP4A3 also decreased the expression of C-terminal Src kinase (CSK, Figure. 6C), which can phosphorylate the tyrosine 527 residue in Src to negatively regulate Src pathway activation [56]. Conversely, PTP4A3 knock-down in Jurkat cells increased total Src protein expression, phosphorylation of Src_Y527, and CSK protein expression (Figure 6E-F). Together, these data show that genetic manipulation of PTP4A3 in T-ALL cells can modulate Src activity by affecting phosphorylation of Src itself as well as affecting expression of kinases known to phosphorylate Src.

In order to further confirm that PTP4A3 affects cell migration via modulation of the Src pathway, we used the Src inhibitor Su6656 to treat PTP4A3 overexpressing cells with or without the PTP4A inhibitor JMS-053 for 2 hours, and evaluated whether the inhibitors synergized to affect cell migration capability in trans-well assays. While both JMS-053 and Su6656 significantly decreased cell migration compared to control, we found no significant additive effect when both inhibitors were used (Figure 6G). These data further support the hypothesis that PTP4A3 modulates Src signaling to promote cell migration.

## 4. Discussion

Treatment outcomes of ALL have been improved dramatically over the last several decades with survival rates exceeding 80%. However, there is a disparity in the number of available immunotherapies and molecular targeted therapies available in T-ALL compared to other leukemia subtypes, correlating with a worse patient prognosis in patients who fail traditional chemotherapy regimens. Additionally, Central Nervous System (CNS) infiltration by T-ALL and both CNS and bone marrow relapse still remains a critical clinical challenge, with survival rates of relapsed disease as low as 40% [57]. The identification and characterization of important drivers of T-ALL progression is needed for the design of novel, targeted therapeutics.

We found that PTP4A3 was highly-expressed in T-ALL patient samples and cell lines compared with healthy PBMCs, which is consistent with studies reporting PTP4A3 upregulation in solid tumors [58] and B-ALL [41]. More importantly, we used two different and unique animal models to demonstrate a oncogenic role for PTP4A3 in T-ALL: overexpression of PTP4A3 in the transgenic zebrafish T-ALL model showed that PTP4A3 expression enhanced spreading of T-ALL cells out of the thymus and promoted their rapid entry into circulation, while human T-ALL cells with silenced PTP4A3 lost their ability to engraft and grow in NSG mice. While this lack of engraftment and growth in mouse xenografts may be attributed to the decrease in migratory ability secondary to the loss of PTP4A3 expression, there may be additional roles that PTP4A3 is playing *in vivo* that partially explain this drastic change in phenotype between PTP4A3 knock-down and control T-ALL cell lines, such as interaction with microenvironment, invasiveness, or other factors. Taken together, these *in vivo* data fortify the critical role of PTP4A3 in T-ALL initiation, migratory spread, and progression.

PTP4A3 knock-down in ovarian cancer [33, 59], gastric cancer [53, 60] and breast cancer [61] impairs cell growth or proliferation, while PTP4A3 overexpression in Human Embryonic Kidney (HEK) [62], AML [17, 26], and ovarian cancer [63] cells increases cell proliferation. Surprisingly, our data shows that T-ALL cell growth was not affected by PTP4A3 knock-down or overexpression. However, cell viability was impaired in cells treated with the pan-PTP4A inhibitor, JMS-053, which might be due to the ability of JMS-053 to also inhibit PTP4A2, which we found to also be highly expressed in T-ALL cells and patient samples and also reported to be important for T-ALL cell proliferation and survival [64].

Our data suggest that a function of PTP4A3 in T-ALL is to promote cell migration. While cell migration is critical to solid tumor progression, and PTP4A3 has well-established roles there, in leukemia, the contribution of migration towards disease progression is less defined. In T-ALL, CCR7, a known regulator of T lymphocyte migration, has been demonstrated to be necessary and sufficient to drive infiltration of T-ALL cells into the CNS in a mouse model [65]. Another regulator of cell migration, CXCR3, has also been reported to be involved in ALL progression, in which CXCR3 inhibition significantly reduced leukemic infiltration into bone marrow, spleen and CNS [66]. Although whether PTP4A3 can modulate known lymphocyte migratory signals, such as CCR7 or CXCR3, to promote T-ALL migration and possibly CNS relapse remains to be determined, there is clearly a precedent for migration playing a critical role in ALL progression.

We have demonstrated that PTP4A3 modulates the Src signaling pathway in T-ALL cell lines. Src, a non-receptor membrane-associated tyrosine kinase, was one of the first discovered oncogenes [67] and its kinase activity is regulated by two critical tyrosine phosphorylation sites, Y416 and Y527. Y527 phosphorylation via CSK leads to an inactive conformation of Src; upon Y527 de-phosphorylation, the inactive conformation is released, leading to auto-phosphorylation of the Y416 residue, which confers maximal kinase activity of Src [56]. Currently it remains unknown which phosphatases are able to specifically regulate Src signaling via dephosphorylation of Y527. Src activation has been reported repeatedly in many types of human cancer, with a prominent role in regulating motility, migration, and metastasis [55]. Moreover, PTP4A3 has been previously reported to play a role in Src pathway activation. For example, endothelial cells isolated from PTP4A3 null mice had decreased Src_pY416 after VEGF stimulation, compared to wild-type cells, resulting in reduced migration and invasiveness [68]. Furthermore, PTP4A3 overexpression in both HEK and colon cancer cells activated the Src signaling pathway via CSK down-regulation, resulting in decreased Src_pY527 and increased cell proliferation and invasion [62]. Similarly, in a recent study of B-ALL, PTP4A3 knock-down abolished the increase in Src_pY416 stimulated by the chemokine, CXCL12, leading to reduced migration and invasion [69].

We have now shown that PTP4A3 modulates the SRC signaling pathway in T-ALL. We found that overexpression of PTP4A3 de-phosphorylates Src_pY527, and that the opposite holds true, with PTP4A3 knock-down leading to an increase in Src_Y527 phosphorylation. However, we also found that PTP4A3 overexpression in T-ALL cell lines resulted in lower CSK protein levels, which conversely increased upon PTP4A3 knock-down. From our data, we cannot conclude whether PTP4A3 acts directly on Src protein, dephophorylating the Y527 residue, or whether it is working via modulation of expression levels of CSK protein, causing an indirect decrease in Src_Y537 phosphorylation. Further studies are necessary to determine if Src is a direct substrate of PTP4A3 in T-ALL.

In summary, we demonstrated an oncogenic role for PTP4A3 in T-ALL, using both *in vitro* and *in vivo* assays. We found PTP4A3 promotes T-ALL development and onset in zebrafish models, and plays a role in engraftment in mouse xenograft models. Cell-culture based assays revealed that PTP4A3 modulates Src signaling in T-ALL to enhance migratory capability, with no significant effect on cell growth. The exact molecular mechanism by which PTP4A3 modulates Src pathway activity needs to be further explored, and we are now in the process of determining the direct substrates of PTP4A3 in T-ALL. Given that several genes involved in T cell migration promote CNS and bone marrow relapse, further studies on the role of PTP4A3 in T-ALL progression, especially in relapse models, are also necessary. There is increasing interest in developing PTP4A3 inhibitors for use in solid tumors; our study indicates that they may be useful in T-ALL as well.

## Abbreviations

T-ALL: T-cell Acute Lymphoblastic Leukemia
B-ALL: B-cell Acute Lymphoblastic Leukemia
AML: Acute Myeloid Leukemia
shRNA: Short-hairpin RNA
PTP4A3: Protein Tyrosine Phosphatase 4A3
CSK: C-terminal Src Kinase
RPPA: Reverse Phase Protein Array
NSG: NOD.Cg-Prkdc^*scid*^Il2rg^tm1Wjl^/SzJ
ANOVA: analysis or variance
FLT3: FMS-like Tyrosine Kinase 3
ITD: Internal Tandem Duplication
AP-1: Activator Protein 1
STR: short tandem repeat
PVDF: polyvinylidene difluoride
HEK: Human Embryonic Kidney
CNS: Central Nervous System

## Acknowledgements

We thank the UK Flow Cytometry & Immune Monitoring Core and the Biostatistics and Bioinformatics Shared Resource Facility (supported by the Markey Cancer Center and an NCI Center Core Support Grant P30CA177558) for assistance with FACS and data analysis, respectively, the MD Anderson RPPA Core facility for assistance with RPPA (funded by P30CA16672) and Hera Biolabs for assistance with mouse xenograft experiments. Donna Gilbreath of the Markey Cancer Center Research Communications Office provided invaluable assistance with graphical editing. We are grateful to John Lazo and Peter Wipf for providing JMS-053, Caroline Smith for synthesizing and providing PTP4A protein, Kristin O’Leary for assistance with zebrafish experiments, and Dylan Rivas for assistance in analyzing patient datasets and histology. This research was supported by a St. Baldrick’s Foundation Research Grant, and NIH grants DP2CA228043, R01CA227656 (to J.S. Blackburn) and NIH Training Grant T32CA165990 (M.G. Haney).

## Author Contributions

M.W. and J.S.B. conceived of and designed the study. M.W. performed all molecular biology and cell-based experiments. M.H. carried out zebrafish experiments. M.W. drafted and edited the manuscript, J.S.B revised. All authors have read and approved of the final version of this manuscript.

## References

1. Van Vlierberghe, P. and A. Ferrando, The molecular basis of T cell acute lymphoblastic leukemia. J Clin Invest, 2012. 122(10): p. 3398–406.

2. Inaba, H., M. Greaves, and C.G. Mullighan, Acute lymphoblastic leukaemia. The Lancet, 2013. 381(9881): p. 1943–1955.

3. Martelli, A.M., et al., Targeting signaling pathways in T-cell acute lymphoblastic leukemia initiating cells. Adv Biol Regul, 2014. 56: p. 6–21.

4. Durinck, K., et al., Novel biological insights in T-cell acute lymphoblastic leukemia. Exp Hematol, 2015. 43(8): p. 625–39.

5. Van Vlierberghe, P., et al., Molecular-genetic insights in paediatric T-cell acute lymphoblastic leukaemia. Br J Haematol, 2008. 143(2): p. 153–68.

6. Bhullar, K.S., et al., Kinase-targeted cancer therapies: progress, challenges and future directions. Molecular cancer, 2018. 17(1): p. 48–48.

7. Zhang, Z.Y., Drugging the Undruggable: Therapeutic Potential of Targeting Protein Tyrosine Phosphatases. Acc Chem Res, 2017. 50(1): p. 122–129.

8. Lazo, J.S. and E.R. Sharlow, Drugging Undruggable Molecular Cancer Targets. Annu Rev Pharmacol Toxicol, 2016. 56: p. 23–40.

9. Ruckert, M.T., et al., Protein tyrosine phosphatases: promising targets in pancreatic ductal adenocarcinoma. Cellular and Molecular Life Sciences, 2019. 76(13): p. 2571–2592.

10. Wei, M., K.V. Korotkov, and J.S. Blackburn, Targeting phosphatases of regenerating liver (PRLs) in cancer. Pharmacology & Therapeutics, 2018. 190: p. 128–138.

11. den Hollander, P., et al., Phosphatase PTP4A3 Promotes Triple-Negative Breast Cancer Growth and Predicts Poor Patient Survival. Cancer Research, 2016. 76(7): p. 1942.

12. Saha, S., et al., A Phosphatase Associated with Metastasis of Colorectal Cancer. Science, 2001. 294(5545): p. 1343.

13. Bardelli, A., et al., PRL-3 Expression in Metastatic Cancers. Clinical Cancer Research, 2003. 9(15): p. 5607–5615.

14. Dai, N., Expression of phosphatase regenerating liver 3 is an independent prognostic indicator for gastric cancer. World Journal of Gastroenterology, 2009. 15(12): p. 1499.

15. Wang, L., et al., PTP4A3 is a target for inhibition of cell proliferatin, migration and invasion through Akt/mTOR signaling pathway in glioblastoma under the regulation of miR-137. Brain Res, 2016. 1646: p. 441–50.

16. Vandsemb, E.N., et al., Phosphatase of regenerating liver 3 (PRL-3) is overexpressed in human prostate cancer tissue and promotes growth and migration. Journal of Translational Medicine, 2016. 14(1): p. 71.

17. Park, J.E., et al., Oncogenic roles of PRL-3 in FLT3-ITD induced acute myeloid leukaemia. EMBO Molecular Medicine, 2013. 5(9): p. 1351.

18. Campbell, A.M. and Z.-Y. Zhang, Phosphatase of regenerating liver: a novel target for cancer therapy. Expert Opinion on Therapeutic Targets, 2014. 18(5): p. 555–569.

19. Yeh, H.C., et al., PTP4A3 Independently Predicts Metastasis and Survival in Upper Tract Urothelial Carcinoma Treated with Radical Nephroureterectomy. J Urol, 2015. 194(5): p. 1449–55.

20. Thura, M., et al., PRL3-zumab as an immunotherapy to inhibit tumors expressing PRL3 oncoprotein. Nature Communications, 2019. 10(1): p. 2484.

21. Radke, I., et al., Expression and prognostic impact of the protein tyrosine phosphatases PRL-1, PRL-2, and PRL-3 in breast cancer. Br J Cancer, 2006. 95(3): p. 347–54.

22. Dai, N., et al., Expression of phosphatase regenerating liver 3 is an independent prognostic indicator for gastric cancer. World Journal of Gastroenterology: WJG, 2009. 15(12): p. 1499–1505.

23. Ren, T., et al., Prognostic significance of phosphatase of regenerating liver-3 expression in ovarian cancer. Pathol Oncol Res, 2009. 15(4): p. 555–60.

24. Mayinuer, A., et al., Upregulation of Protein Tyrosine Phosphatase Type IVA Member 3 (PTP4A3/PRL-3) is Associated with Tumor Differentiation and a Poor Prognosis in Human Hepatocellular Carcinoma. Annals of Surgical Oncology, 2013. 20(1): p. 305–317.

25. Beekman, R., et al., Retroviral integration mutagenesis in mice and comparative analysis in human AML identify reduced PTP4A3 expression as a prognostic indicator. PLoS One, 2011. 6(10): p. e26537.

26. Qu, S., et al., Independent oncogenic and therapeutic significance of phosphatase PRL-3 in FLT3-ITD-negative acute myeloid leukemia. Cancer, 2014. 120(14): p. 2130–41.

27. Zeng, Q., et al., PRL-3 and PRL-1 Promote Cell Migration, Invasion, and Metastasis. Cancer Research, 2003. 63(11): p. 2716.

28. Guo, K., et al., Catalytic domain of PRL-3 plays an essential role in tumor metastasis: Formation of PRL-3 tumors inside the blood vessels. Cancer Biology & Therapy, 2004. 3(10): p. 945–951.

29. Wu, X., et al., Phosphatase of Regenerating Liver-3 Promotes Motility and Metastasis of Mouse Melanoma Cells. The American Journal of Pathology, 2004. 164(6): p. 2039–2054.

30. Hardy, S., et al., Overexpression of the Protein Tyrosine Phosphatase PRL-2 Correlates with Breast Tumor Formation and Progression. Cancer Research, 2010. 70(21): p. 8959.

31. Kato, H., et al., High expression of PRL-3 promotes cancer cell motility and liver metastasis in human colorectal cancer. A Predictive Molecular Marker of Metachronous Liver and Lung Metastases, 2004. 10(21): p. 7318–7328.

32. Li, Z., et al., Inhibition of PRL-3 gene expression in gastric cancer cell line SGC7901 via microRNA suppressed reduces peritoneal metastasis. Biochemical and Biophysical Research Communications, 2006. 348(1): p. 229–237.

33. Polato, F., et al., PRL-3 Phosphatase Is Implicated in Ovarian Cancer Growth. Clinical Cancer Research, 2005. 11(19): p. 6835–6839.

34. Qian, F., et al., PRL-3 siRNA Inhibits the Metastasis of B16-BL6 Mouse Melanoma Cells In Vitro and In Vivo. Molecular Medicine, 2007. 13(3-4): p. 151–159.

35. Zimmerman, M.W., G.E. Homanics, and J.S. Lazo, Targeted deletion of the metastasis-associated phosphatase Ptp4a3 (PRL-3) suppresses murine colon cancer. PLoS One, 2013. 8(3): p. e58300.

36. McQueeney, K.E., et al., Targeting ovarian cancer and endothelium with an allosteric PTP4A3 phosphatase inhibitor. Oncotarget, 2018. 9(9): p. 8223–8240.

37. Bai, Y., et al., Novel Anticancer Agents Based on Targeting the Trimer Interface of the PRL Phosphatase. Cancer Res, 2016. 76(16): p. 4805–15.

38. Hoeger, B., et al., Biochemical evaluation of virtual screening methods reveals a cell-active inhibitor of the cancer-promoting phosphatases of regenerating liver. European Journal of Medicinal Chemistry, 2014. 88: p. 89–100.

39. Thura, M., et al., PRL3-zumab, a first-in-class humanized antibody for cancer therapy. JCI Insight, 2016. 1(9): p. e87607.

40. Zhou, J., et al., PRL-3, a metastasis associated tyrosine phosphatase, is involved in FLT3-ITD signaling and implicated in anti-AML therapy. PLoS One, 2011. 6.

41. Hjort, M.A., et al., Phosphatase of regenerating liver-3 is expressed in acute lymphoblastic leukemia and mediates leukemic cell adhesion, migration and drug resistance. Oncotarget, 2017. 9(3): p. 3549–3561.

42. Langenau, D.M., et al., Myc-Induced T Cell Leukemia in Transgenic Zebrafish. Science, 2003. 299(5608): p. 887–890.

43. Blackburn, J.S., et al., Notch signaling expands a pre-malignant pool of T-cell acute lymphoblastic leukemia clones without affecting leukemia-propagating cell frequency. Leukemia, 2012. 26(9): p. 2069–2078.

44. Gürtler, A., et al., Stain-Free technology as a normalization tool in Western blot analysis. Analytical Biochemistry, 2013. 433(2): p. 105–111.

45. Gallo, A., et al., Gross Cystic Disease Fluid Protein-15(GCDFP-15)/Prolactin-Inducible Protein (PIP) as Functional Salivary Biomarker for Primary Sjögren’s Syndrome. Journal of genetic syndromes & gene therapy, 2013. 4: p. 10.4172/2157-7412.1000140.

46. Kohlmann, A., et al., An international standardization programme towards the application of gene expression profiling in routine leukaemia diagnostics: the Microarray Innovations in LEukemia study prephase. British journal of haematology, 2008. 142(5): p. 802–807.

47. Haferlach, T., et al., Clinical utility of microarray-based gene expression profiling in the diagnosis and subclassification of leukemia: report from the International Microarray Innovations in Leukemia Study Group. Journal of clinical oncology: official journal of the American Society of Clinical Oncology, 2010. 28(15): p. 2529–2537.

48. Winter, S.S., et al., Identification of genomic classifiers that distinguish induction failure in T-lineage acute lymphoblastic leukemia: a report from the Children’s Oncology Group. Blood, 2007. 110(5): p. 1429.

49. Gutierrez, A., et al., The BCL11B tumor suppressor is mutated across the major molecular subtypes of T-cell acute lymphoblastic leukemia. Blood, 2011. 118(15): p. 4169.

50. Tibes, R., et al., Reverse phase protein array: validation of a novel proteomic technology and utility for analysis of primary leukemia specimens and hematopoietic stem cells. Molecular Cancer Therapeutics, 2006. 5(10): p. 2512–2521.

51. Blackburn, J., Liu, Sali, Langenau, David M., Quantifying the Frequency of Tumor-propagating Cells Using Limiting Dilution Cell Transplantation in Syngeneic Zebrafish. JoVE, 2011(53): p. e2790.

52. Milne, S.B., et al., A targeted mass spectrometric analysis of phosphatidylinositol phosphate species. J Lipid Res, 2005. 46(8): p. 1796–802.

53. Matsukawa, Y., et al., Constitutive Suppression of PRL-3 Inhibits Invasion and Proliferation of Gastric Cancer Cell in vitro and in vivo. Pathobiology, 2010. 77(3): p. 155–162.

54. Lin, M.-D., et al., Expression of phosphatase of regenerating liver family genes during embryogenesis: an evolutionary developmental analysis among Drosophila, amphioxus, and zebrafish. BMC Developmental Biology, 2013. 13(1): p. 18.

55. Guarino, M., Src signaling in cancer invasion. Journal of Cellular Physiology, 2010. 223(1): p. 14–26.

56. Okada, M., Regulation of the SRC family kinases by Csk. International journal of biological sciences, 2012. 8(10): p. 1385–1397.

57. Cannon, J.L., S.R. Oruganti, and D.W. Vidrine, Molecular regulation of T-ALL cell infiltration into the CNS. Oncotarget, 2017. 8(49): p. 84626–84627.

58. Bollu, L.R., et al., Molecular Pathways: Targeting Protein Tyrosine Phosphatases in Cancer. Clinical Cancer Research, 2017. 23(9): p. 2136–2142.

59. Matter, W.F., et al., Role of PRL-3, a human muscle-specific tyrosine phosphatase, in angiotensin-II signaling. Biochem Biophys Res Commun, 2001. 283(5): p. 1061–8.

60. Wang, Z., et al., High Expression of PRL-3 can Promote Growth of Gastric Cancer and Exhibits a Poor Prognostic Impact on Patients. Annals of Surgical Oncology, 2009. 16(1): p. 208–219.

61. Gari, H.H., et al., Loss of the oncogenic phosphatase PRL-3 promotes a TNF-R1 feedback loop that mediates triple-negative breast cancer growth. Oncogenesis, 2016. 5(8): p. e255.

62. Liang, F., et al., PRL3 Promotes Cell Invasion and Proliferation by Down-regulation of Csk Leading to Src Activation. Journal of Biological Chemistry, 2007. 282(8): p. 5413–5419.

63. Huang, Y.H., et al., A role of autophagy in PTP4A3-driven cancer progression. Autophagy, 2014. 10(10): p. 1787–800.

64. Kobayashi, M., et al., Phosphatase PRL2 promotes oncogenic NOTCH1-Induced T-cell leukemia. Leukemia, 2017. 31(3): p. 751–754.

65. Buonamici, S., et al., CCR7 signalling as an essential regulator of CNS infiltration in T-cell leukaemia. Nature, 2009. 459(7249): p. 1000–1004.

66. Gómez, A.M., et al., Chemokines and relapses in childhood acute lymphoblastic leukemia: A role in migration and in resistance to antileukemic drugs. Blood Cells, Molecules, and Diseases, 2015. 55(3): p. 220–227.

67. Martin, G.S., TIMELINE: The hunting of the Src. Nature Reviews Molecular Cell Biology, 2001. 2(6): p. 467–475.

68. Zimmerman, M.W., et al., Protein-tyrosine phosphatase 4A3 (PTP4A3) promotes vascular endothelial growth factor signaling and enables endothelial cell motility. J Biol Chem, 2014. 289(9): p. 5904–13.

69. Hjort, M.A., et al., Phosphatase of regenerating liver-3 (PRL-3) is overexpressed in classical Hodgkin lymphoma and promotes survival and migration. Experimental Hematology & Oncology, 2018. 7(1): p. 8.

