## Supplemental Table 1 for "Protein Tyrosine Phosphatase 4A3 (PTP4A3) modulates Src signaling in T-cell Acute Lymphoblastic Leukemia to promote leukemia migration and progression"

**Table S1: List of antibodies, their source and working dilutions**

| **Reagents** | **Source** | **Host species** | **Working dilution** |
| --- | --- | --- | --- |
| Anti-PTP4A3 | Abcam (ab50276, Lot GR9977-20) | Rabbit | 1:500 |
| Anti-PTP4A | R&D(MAB32191, Lot XJS01 ) | Mouse | 1:1000 |
| Anti-Src | Cell Signaling (Clone 32G6, 2123, Lot 5) | Rabbit | 1:1000 |
| Anti-Src_pY416 | Cell Signaling (Clone D49G4, 2123, Lot 4) | Rabbit | 1:1000 |
| Anti-Src_pY527 | Cell Signaling (2105, Lot 9) | Rabbit | 1:1000 |
| Anti-CSK | Cell Signaling (Clone C74C1,4980, Lot 2) | Rabbit | 1:1000 |
| PE Anti-Human CD45 | Biolegend (Clone HI30, 304008) | Mouse | 1:1000 |
| Mouse IgG-HRP | Cell Signaling (7106, Lot TC2625) | Goat | 1:5000 |
| Rabbit IgG-HRP | GeneTex (26741, Lot 9788061) | Goat | 1:5000 |
| Blocking buffer | 5% milk in 1% TBST |  |  |
