## Supplemental Figures 1-5 for "Protein Tyrosine Phosphatase 4A3 (PTP4A3) modulates Src signaling in T-cell Acute Lymphoblastic Leukemia to promote leukemia migration and progression"

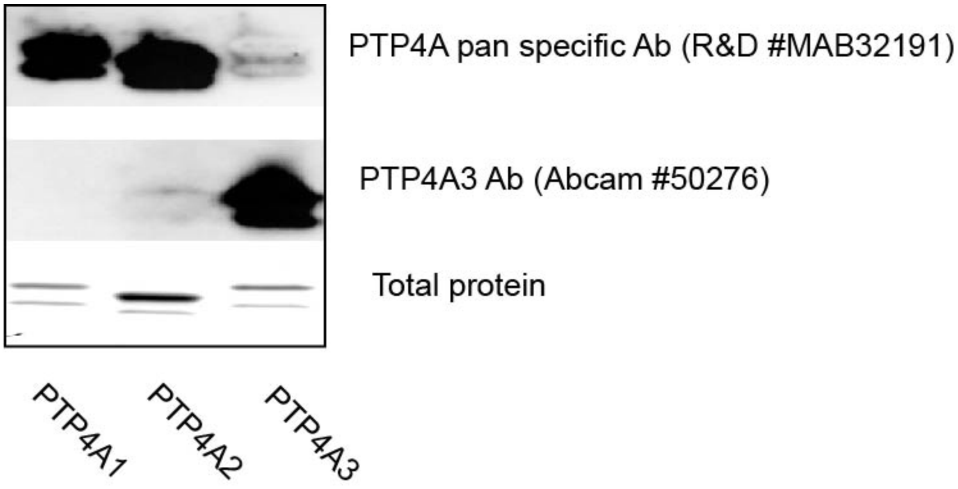


**Supplemental Figure 1. PTP4A antibody validation.** Antibody against PTP4A3 (Abcam, 50276) detects purified PTP4A3 protein specifically but not PTP4A1 and 2. Human PTP4A pan specific antibody (MAB32191) detects both PTP4A1 and PTP4A2 at similar sensitivity but barely detects PTP4A3. Recombinant human PTP4A1 (8490-PT), PTP4A2 (6694-PT) and PTP4A3 (8455-PT) proteins were from R&D systems and 0.3 µg of each protein was loaded in 4-20% protein TGX stain-free gel (BioRad, cat #4568093).

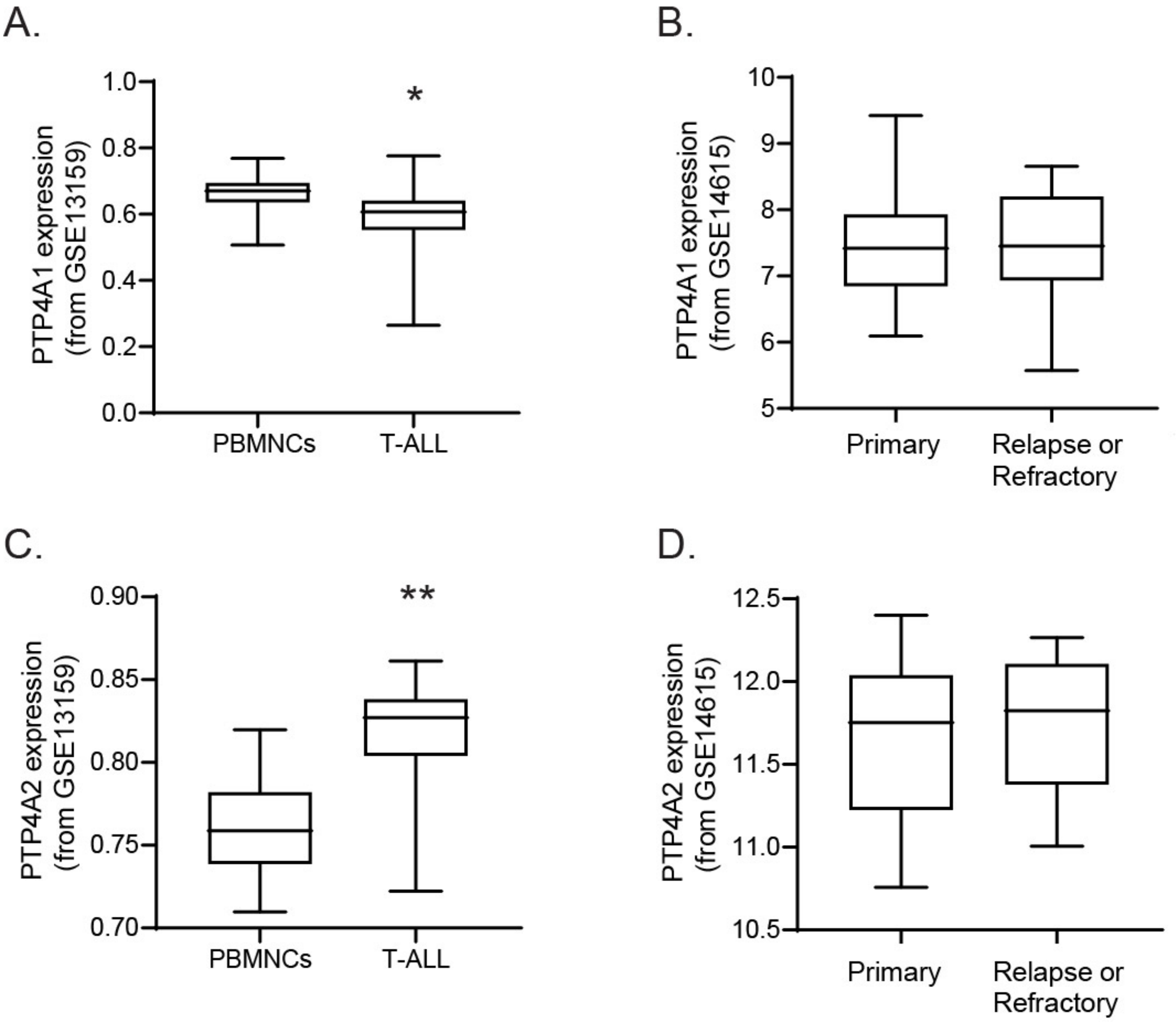


**Supplemental Figure 2. Expression of PTP4A1 and PTP4A2 in T-ALL.** Microarray expression analysis of PTP4A1 (A) and PTP4A2 (C) expression in primary T-ALL compared to peripheral blood mononuclear cells (PBMCs), and PTP4A1 (B) and PTP4A2 (D) in primary T-ALL compared to relapsed and refractory disease *$p<0.0001$ PTP4A1 in T-ALL compared to PBMNC, **$p<0.001$ PTP4A2 expression in T-ALL compared to PMBNC.

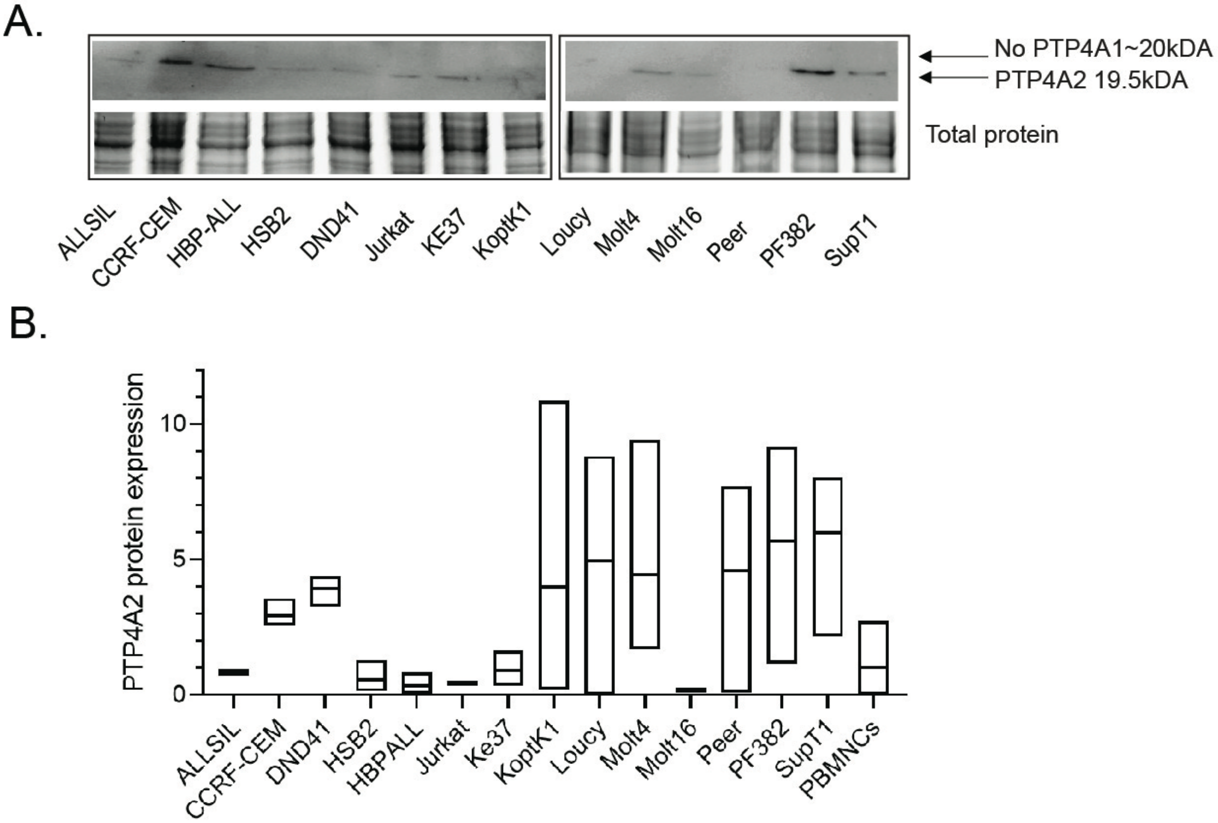


**Supplemental Figure 3. T-ALL cells lines express PTP4A2 but not PTP4A1.** (A) Representative western blot analysis of PTP4A1 and PTP4A2 in human T-ALL cell lines, showing PTP4A2 expression in a subset of T-ALL cell lines. (B) Quantification of 3 independent western blots cross human T-ALL cell lines, showing that PTP4A2 expression can vary, even with the same cell line.

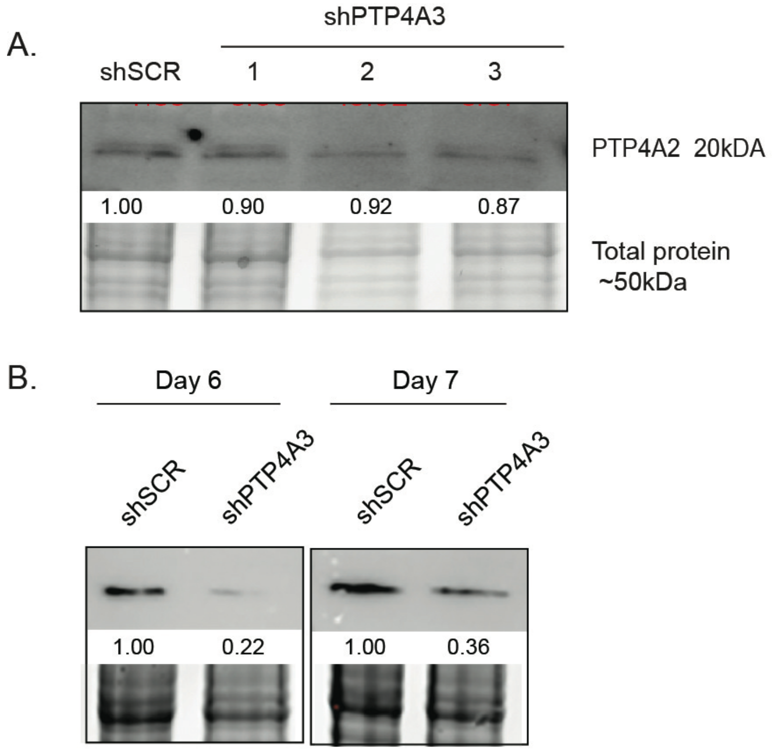


**Supplemental Figure 4. Effects of PTP4A3 knock-down.** Western blot analysis shows PTP4A2 expression does not change upon PTP4A3 knock-down, indicating PTP4A2 does not compensate for PTP4A3 loss. (B) Jurkat cells, which showed PTP4A3 complete knock-down at day 5 post lentiviral infection (Figure 3) were cultured for 6 and 7 days and total protein was extracted for western blot analysis, showing that PTP4A3 expression returns over time. Blots are representative of at least 3 independent experiments.

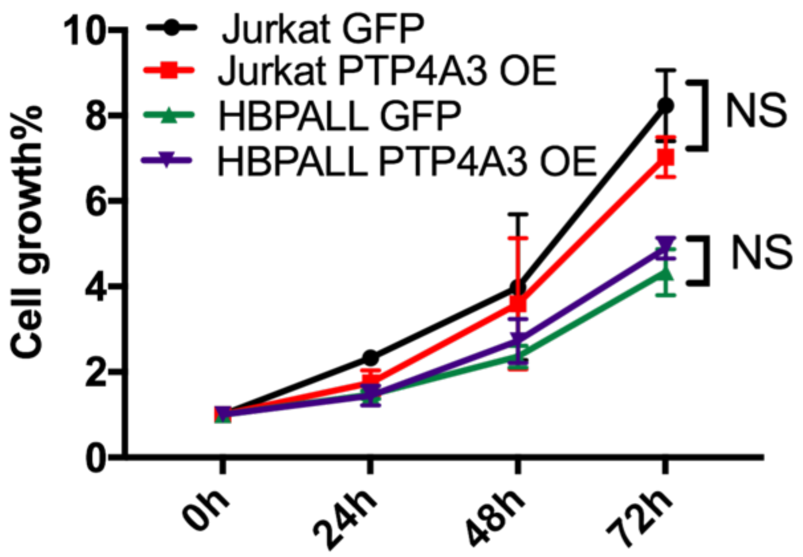


**Supplemental Figure 5. PTP4A3 overexpression does not impact growth of Jurkat or HBPALL cells.** T-ALL cells expressing PGK:PTP4A3 (PTP4A3 OE) or PGK:GFP (GFP) were cultured in medium with puromycin for 72 hours. Cell growth was determined by Cell Titer-glow assay and normalized to the readout of day 0, and showed no difference between PTP4A3 overexpressing cells and control cells. Data shown are the average of 3 independent experiments, done in triplicate, NS= not significant.
